# An alternative class of targets for microRNAs containing CG dinucleotide

**DOI:** 10.1101/026708

**Authors:** Zheng Yan, Bin Zhang, Haiyang Hu, Gangcai Xie, Song Guo, Philipp Khaitovich, Yi-Ping Phoebe Chen

**Author notes:** These authors contributed equally to this work (Zheng Yan, Bin Zhang, Haiyang Hu). Corresponding author Email addresses.

## Abstract

**Background:** MicroRNAs are endogenous ∼23nt RNAs which regulate mRNA targets mainly through perfect pairing with their seed region (positions 2-7). Several instances of bulge UTR sequence can also be recognized by miRNA as their target. But such non-Watson-Crick base pairings are incompletely understood.

**Results:** We found a group of miRNAs which had very few conservative targets while potentially having a subclass of bulge message RNA targets. Compared with the canonical target, these bulge targets had a lower negative correlation with the miRNA expression, and either were downregulated in the miRNA overexpression experiment or upregulated in the miRNA knock-down experiment.

**Conclusions:** We proved that the bulge target exists widely in certain groups of miRNAs and such non-canonical targets can be recoginized by miRNA. Incorporating these bulge targets, combined with evolutionary conservation, will reduce the false-positive rate of microRNA computational target prediction.

## Background

MicroRNAs(miRNAs) are ∼23 nucleotide RNAs that regulate eukaryotic gene expression post-transcriptally [1]. miRNAs use base-pairing to guide RNA-induced silencing complexes (RISCs) to specific message RNAs with fully or partly complementary sequences, primarily in the 3’ untranslated region [2]. The best characterized features determining animal miRNA-target recognition are six-nucleotide (nt) long seed sites, which perfectly complement the 5’ end of the miRNA (positions 2-7) [3]. This Watson-Crick seed pairing rule is sufficient on its own for predicting conserved targets above the noise of false-positive predictions in most miRNAs [4].

Most of the miRNA-target prediction algrorithms rely heavily on seed rules and evolutionary conservation [5, 6]. However, such strategies suffer from missing the noncanonical target sites [7]. Several bioloical studies have functionally validated the existence of imperfect binding sites [8–10].

Recently, Ago HITS-CLIP was used to precisely map the miRNA-binding sites in both *Caenorhabditis elegans* [11] and mouse brains [7]. However, about one-quarter of the total binding sites did not follow the classical seed rules in mouse brains [7]. Further analysis revealed that the miR-124, one of the most abundant miRNAs in Ago complex in mouse brains, has plenty of noncanonical bulge sites. More recently, an improved CLIP-seq method, CLASH (cross linking, ligation and sequencing of hybrids), revealed around 60% of the seed interactions are noncanonical, containing bulged or mismatched nucleotides [12].

Although these studies strongly suggest the existence of bulge sites, the general features of their interactions with miRNAs are largely unknown, partly due to the difficulty in determining how frequently such atypical sites are used in vivo and what are the general rules to predict them.

Here, we analyze a group of highly conserved miRNAs in verterbrate, but with relative fewer conservative target using the seed rule. Meanwhile, these miRNAs all have a common feature, this being that their seed region contains CG dinucleotide (hereafter refer as CG dimer). We found these potential miRNA regulatory sites have a nucleotide bulge compared with a fully complementary sequence. This expands our insight into miRNA-target interaction.

## Results

### MicroRNA containing CG dimer has fewer cononical targets

Evolutionary conservation has been widely used to identify miRNA-binding sites together with the seed rule. We searched for the orthologs of all the miRNAs annotated by miRbase (miRbase version 17) [13] using their mature sequence in the genomes of 23 species (Supplement Table 1). 1426 annotated miRNAs were divided into three categories, verterbrate, mammal and primate conservative miRNA. TargetScan [14] was used to look for the verterbrate conservative miRNAs’ cononical targets. According to the target site conservative value (Table 1), a class of miRNAs containing CG dimer in their seed region have both very few cononical targets (t test, p < 0.01) and much fewer conservative target sites than the rest of the miRNAs (t test, p < 0.01). For the mammal and primate conservative miRNAs, miRNAs with CG dimer in their seed region also have much fewer conservative target sites (Supplementary Table 2, 3) (t test, p < 0.01).

### Identification of bulge sites that pair to microRNA containing CG dimer

To uncover the possible bulge site, we allow one nucleotide insertion in every position in the seed region (Figure 1) for all the verterbrate conservative miRNAs. Using these artificial seed sequences, we find that only the bulge site inserted between CG dimer can increase the target number and conservation of the target sites (Figure 2). In contrast, the random bulge at the target binding site did not increase the conservation rate.

**Figure 1.**
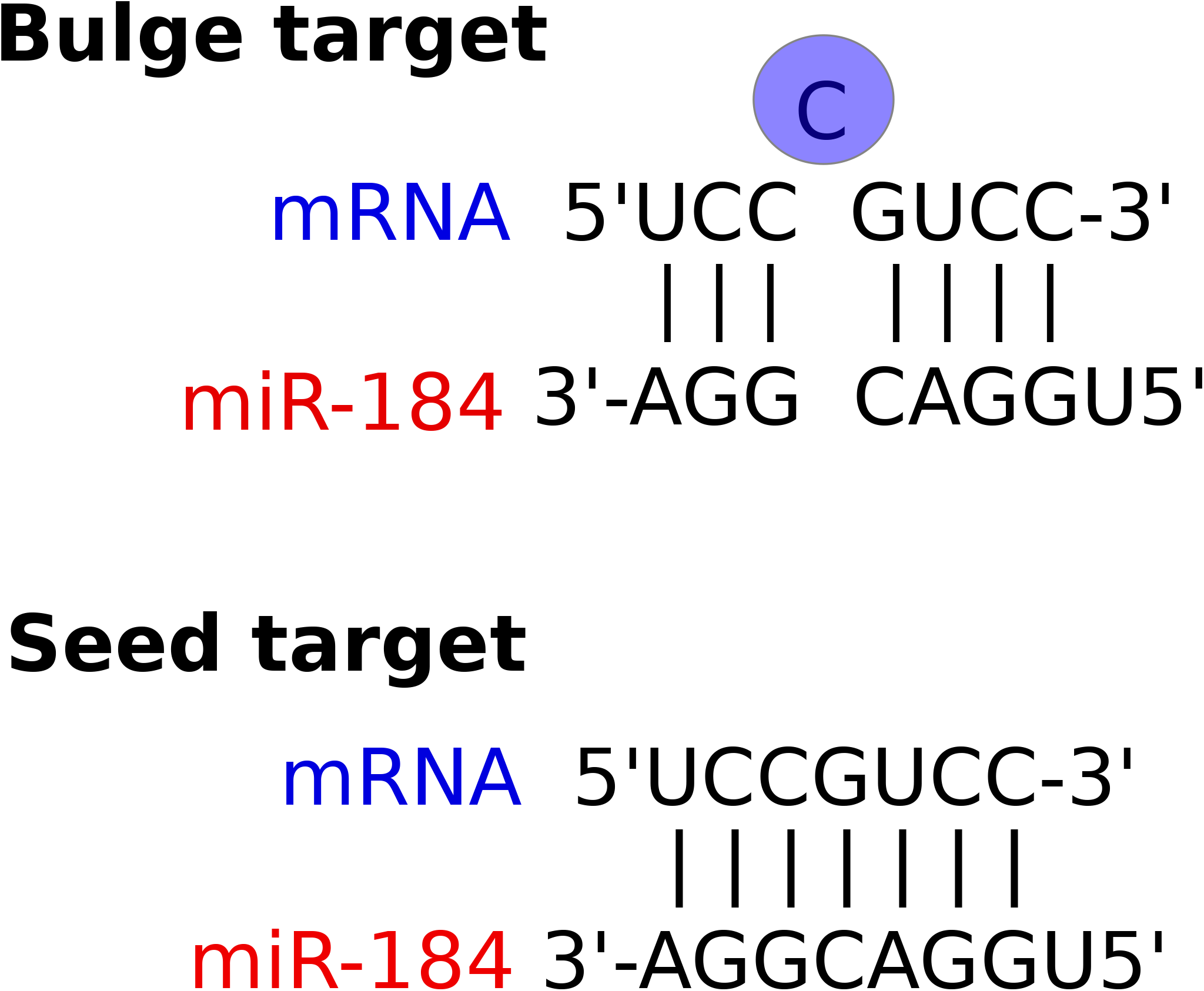
miRNA and Seed/bulge target duplex model. The miR-184 seed sequence was used to illustrate a canonical target match, and a non-canonical target match with a bulge nuleotide between the CG dinucleotide.

**Figure 2.**
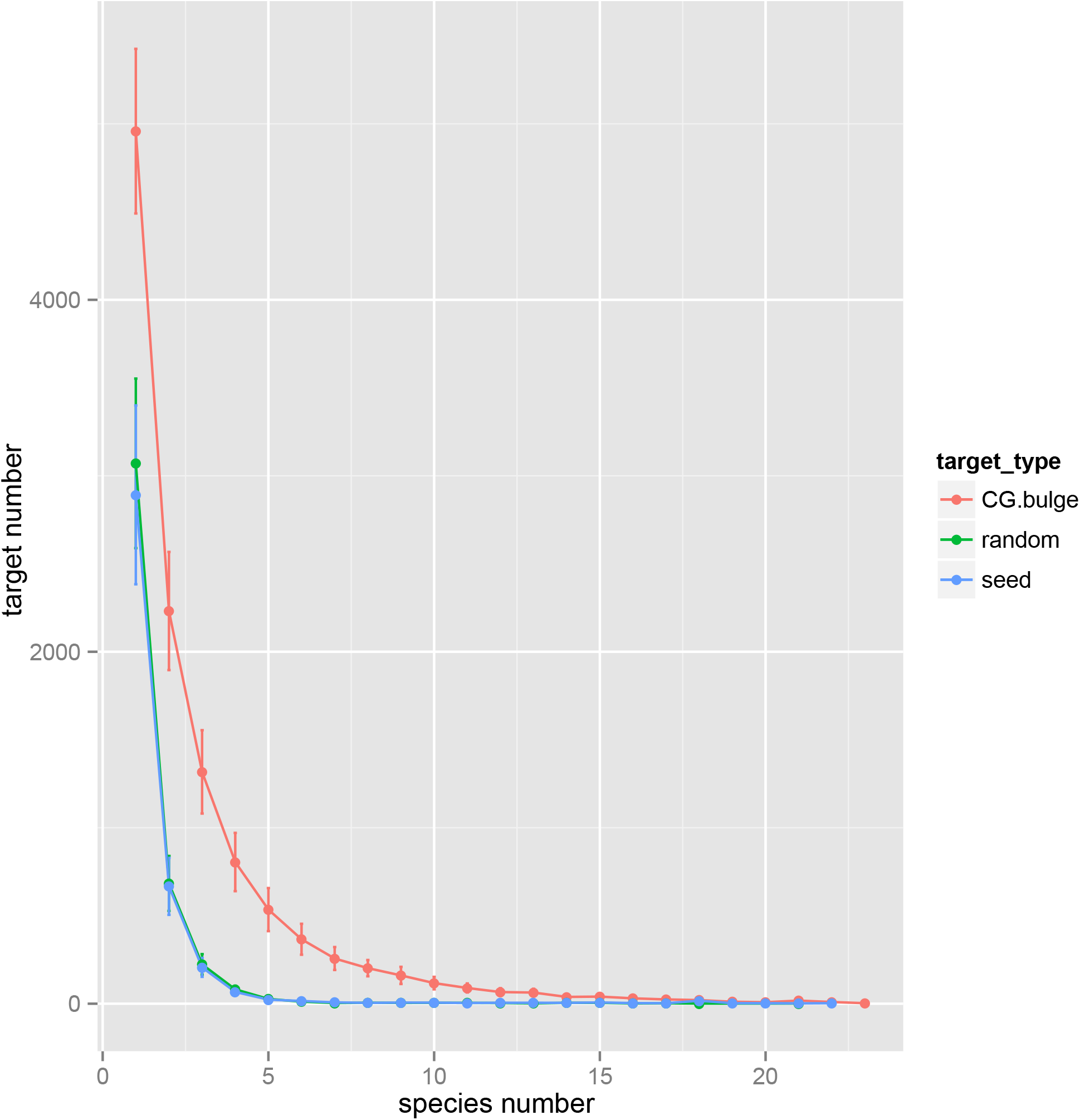
Targets with Bulge between CG dimer have higher conservative rate. Two different target site conservation rates in 23 vertebrate compared with a random bulge target.

### Transcriptome-wide evidence for miRNA repression through bulge target site

We used human age series mRNA and miRNA expression data [15] to quantify the transcriptome correlation between CG dimer miRNAs and their bulge target. The bulge target genes are significantly more negatively correlated to their miRNAs’ expression than the background (Wilcox test, p < 0.01) and for miR-191, the bulge target even outperforms the seed target (Wilcox test, p < 0.01) (Figure 3). We also use public data on transcriptome change after over-expression or knockdown individual miRNAs from GEO. For miR-126, miR210 and miR-184, all the bulge targets were significantly down-regulated after over-expression (Table 2) and in the case of the knock-down experiment for miR-1204, the bulge target is also much more highly expressed compared with the non-target gene (Wilcox test, p < 0.01).

**Figure 3.**
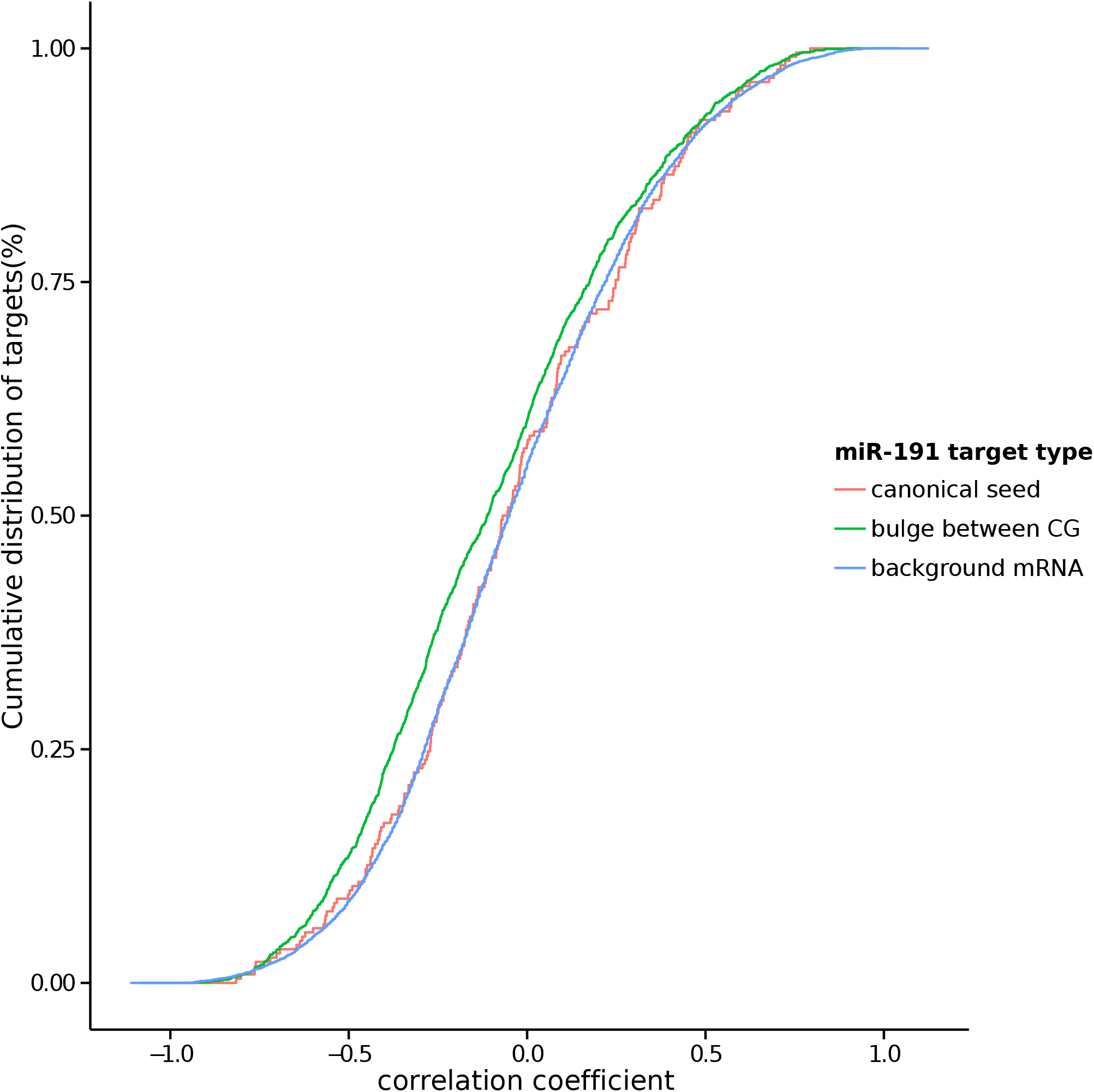
CG dimer miRNA suppression of the bulge target expression. Cumulative distribution of correlation coefficient between miR-191 and target expression level.

### Free energies of CG bulge target duplexes are significantly lower than the random bulges

We compared the mimuim free energy (MFE) between the canonical target, CG bulge target and target with random bulges using RNAhybrid [16]. The non-canonical target with a bulge between CG has a significantly lower MFE compared with the target with random bulge (Wilcox test, p < 0.05, Figure 4).

**Figure 4.**
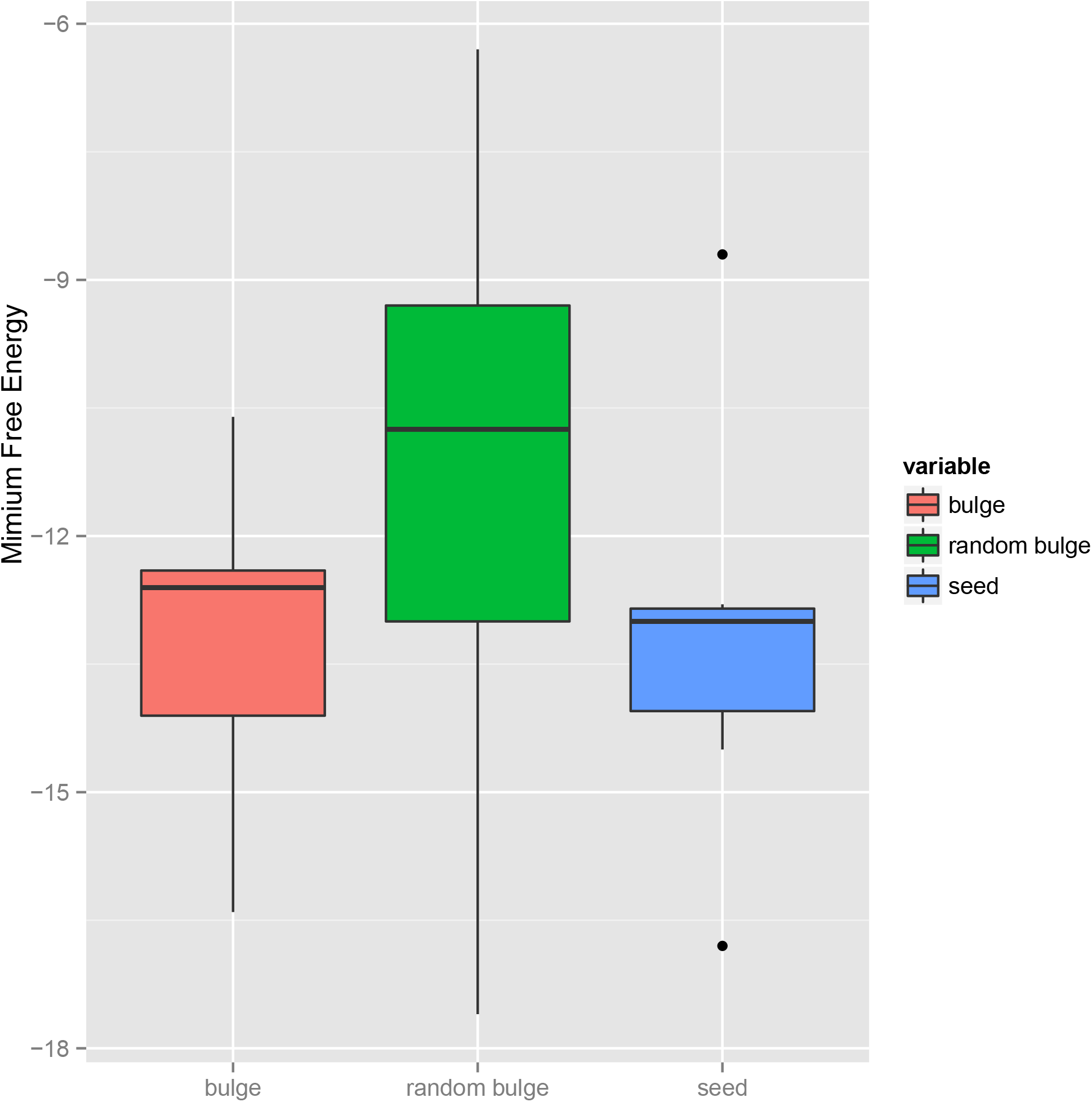
Mimium Free energy between seed pair, CG bulge and Random bulge. Boxplot of the distrubution of the canonical target, noncanonical bulge interaction and random bulge target.

### Validation of the bulge target site by the CLASH data

To allow direct mapping of miRNA-target interactions, we use the CLASH dataset [12] to validate our bulge target for the miRNAs containing CG dimer. Briefly, the RNA molecules present in AGO-associated miRNA-target duplexes were partially hydrolyzed, ligated, reverse transcribed and subjected to illumina sequencing. Compared with the HITS-CLIP and PAR-CLIP dataset, CLASH technology generated a group of reads which contain the miRNAs and their target site sequence together (chimeric reads). In all the six independent CLASH experiments, we find 10 CG dimer miRNAs were detected in all the chimeric reads and 8 miRNAs had, in total, 264 chimeric reads containing a bulge nucleotide between the CG dinucleotide target site (Supplementary Table 4). For all miRNAs detected in the CLASH dataset, the non-canonical interactions (G.U pairs, all possible one nt mismatch or bulge; non-canonical seed) were about 1.7-fold more than the perfect seed base pairing. But within the CG miRNA, only the bulge targets between CG dimer, which in comparison to randomized sequences, showed strong enrichment among all the interactions (Figure 5).

**Figure 5.**
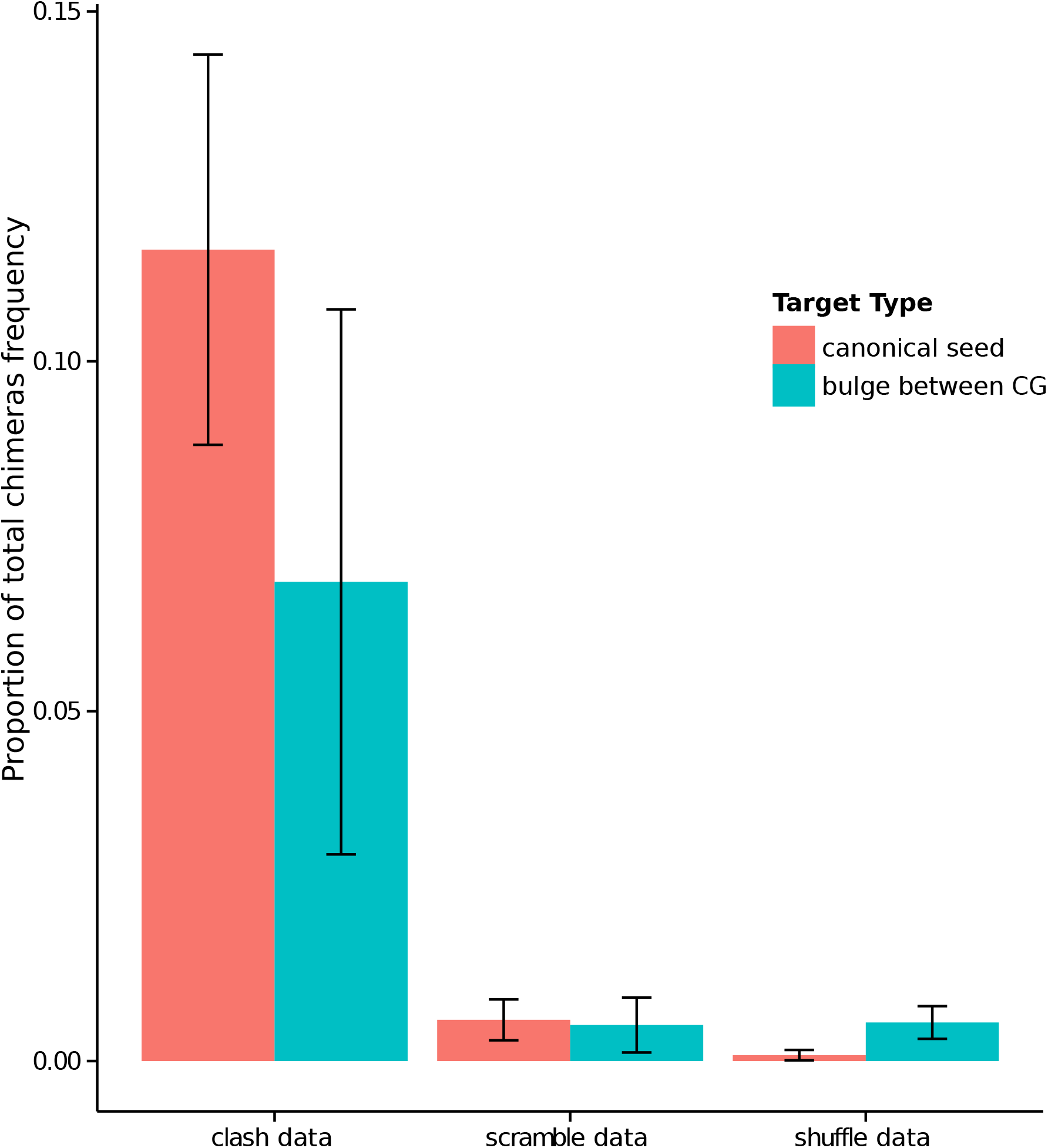
CLASH data validation o f CpG containing miRNA and bulge target interaction. Proportion of canonical seed interaction, noncanonical bulge interaction among CLASH chimeras, scramble and shuffle data.

## Discussion

The aim of this study is to identify the general features of miRNAs and their bulge target interactions. We used the non-canonical miRNA target’s interactome which contains bulge nucleotide between CpG dinucleotide to test whether a bulge position is random or has specific rules. This sub-class of miRNA was observed to have a few seed targets and these seed targets are evolutionally less conservative, which makes these miRNAs potantially non-canonical target rich. Multistep validation, which included evolutionary, overexpression, correlation and CLASH analysis, supports the reliability that between CpG in the seed region, there is a bulge containing a target group.

Multiple studies have found the existence of a bulge target for miRNAs. In particular, the PAR-CLIP and HIS-CLIP technique proved the abundance of non-canonical targets binding to the RISC complex, and the bulge target was consided as one of the major types within such a non-canonical target. However, the short of high-throughput way to detect miRNA-target duplex makes it difficult to predict the position of the hot spot of the bulge nucleotide. Thus, most computational algorithms are restricted to predict only the seed target.

Notably, different miRNAs vary in the target interaction pattern. A previous study [17] also showed that the composition of seed sequences is a major determinant of the miRNA target pattern. However, of all the different kinds of non-canonical target groups, even with the CLASH dataset, there are still no key features to distinguish any miRNA group with a higher bulge target proportion.

Intriguingly, the CG bulge target genes are functionally enriched in synapse and neruon projection and development, indicating that fast evolved neuron cells are most sensitive to the pertubation of seed complementary base-pairing and most receptive in accomodating evolutionary innovation, such as bulge target recognition.

Thus, a major novelty of this work is the identification of a sequence motif, CG dimer, in the seed region of miRNAs is strongly correlated to bulge targeting patterns. This seed variability issue should be taken into consideration when predicting targets for individual miRNAs.

Overall, we found that the bulge targets were preferentially associated with the miRNAs containing CG dinucleotide in their seed region.

## Methods

### miRNA sequences and 3′UTR sequence alignments

Mature miRNA sequences were obtained from the miRBase website (http://www.mirbase.org) [13]. miRNAs are considered to be conserved if they share the same mature sequence in different groups of species, vertebrate, mammal and primates. Genomic coordinates of Ensembl human genes (hg18) were used to extract the human 3′UTR sequences and the corresponding aligned sequences from the 28-species alignment (MAF file) available at the UCSC Table browser. Only protein coding genes were included in the database and when several mRNA isoforms were reported for the same Ensembl gene ID, only the one with the longest 3′UTR sequence was used in the analysis.

### Predictions of seed and bulge target for conservative CG dimer miRNAs

The seed sequences for the CG dimer miRNAs were extracted to find three types of targets. Any coding gene’s 3’UTR containing a perfect complementary sequence was defined as a seed target. For the bulge target, we allowed one extra nucleotide to exist between CG dimer. Randomly inserted single nucleotide seed sequences were used as control. The occurences of the homologous target sites in different species were summed up for seed, bulge and control separately as the conservation rates.

### miRNAs and target expression correlation analysis

12 human brain prefrontal cortex samples’ miRNA (GSE29356) and coding gene transcriptome data (GSE22570) were used to check expression correlation. Both the Pearson and Spearman methods were used to calculate the correlation. For miRNA overexprssion in vitro, miR-126 in LM2 breast cancer cell (GSE23905), miR-184 in SY5Y (GSE26545) and miR-210 in MCF-7 cells and MDA-MB-231.cells (GSE25162) were downloaded from GEO and for the miRNA knockdown experiment, miR-126 in MDA-MB-231 cells and miR-1204 in SUM159PT (GSE37185). The Mann-Whitney-Wilcoxon Test was used to test the seed and bulge target expression change in the transfection experiment.

### Confirm bulge target with CLASH dataset

The miRNA-mRNA interaction sequences was download from the joural’s website in the supplementary Data section [12]. In this published CLASH dataset, the crosslinked RNA-induced silencing complex (RISC) in HEK293 cells were immunoprecipitated. The miRNA and cognate mRNA target transcripts were ligated and sequenced together. The chimeric reads containing vertebrate conservative CG dimer miRNAs were extracted. The bulge target was recognized if there is one extra nucleotide between CG dimer.

### Authors’ contributions

Z.Y and B.Z performed the computational analyses. Y.P.C and Z.Y designed the research and wrote the paper.

## Acknowledgements

We thank the members of the Khaitovich and Chen laboratories for helpful discussions. We thank Xiaowei Wang for the suggestion. This work was supported by La Trobe University Postgraduate Scholarship.

## Tables

**Table 1 - Vertebrate conservative miRNA TargetScan result** All the conservative miRNAs in 23 vertebrate species listed in Supplementary Table 1 were used to calulate their canonical target conservative rates by TargetScan. The results were sorted from lowest conservative rate to highest.

**Table 2 - Functional analysis of CpG miRNA and their bulge targets** The transcriptome data after overexpression or knockdown studies. The Mann-Whitney-Wilcoxon Test was used to test the seed and bulge target expression change compared with random genes.

## Additional files

**Supplementary Table 1 – All the 23 species which were used for miRNA and target analysis.**

All the miRNAs and their target conservation scores were calculated based on these 23 species. File was in Excel format.

**Supplementary Table 2 – Mammal conservative miRNA TargetScan result**

All the miRNAs which have same mature sequence in the mammal species were tested were sorted according to their target site conservation rate. File was in Excel format.

**Supplementary Table 3 – Primate conservative miRNA TargetScan result**

All the miRNAs which have same mature sequence in the primate species we tested were sorted according their target site conservation rate. File was in Excel format.

**Supplementary Table 4 – CLASH chimeras reads which the miRNA-target duplex were sequenced**

All the CG miRNAs which were detected together with their bulge targets between CG dimer are listed in this table. File was in Text format.

